# Resolving the Structure of a Guanine Quadruplex in TMPRSS2 Messenger RNA by Circular Dichroism and Molecular Modeling

**DOI:** 10.1101/2024.07.29.605618

**Authors:** Luisa D’Anna, Aurane Froux, Aurianne Rainot, Angelo Spinello, Ugo Perricone, Florent Barbault, Stéphanie Grandemange, Giampaolo Barone, Alessio Terenzi, Antonio Monari

## Abstract

The presence of a guanine quadruplex in the opening reading frame of the messenger RNA coding for the transmembrane serine protease 2 (TMPRSS2) may pave the way to original anticancer and host-oriented antiviral strategy. Indeed, TMPRSS2 in addition to being overexpressed in different cancer types, is also related to the infection of respiratory viruses, including SARS-CoV-2, by promoting the cellular and viral membrane fusion through its proteolytic activity. The design of selective ligands targeting TMPRSS2 messenger RNA requires a detailed knowledge, at atomic level, of its structure. Therefore, we have used an original experimental-computational protocol to predict the first resolved structure of the parallel guanine quadruplex secondary structure in the RNA of TMPRSS2, which shows a rigid core flanked by a flexible loop. This represents the first atomic scale structure of the guanine quadruplex structure present in TMPRSS2 messenger RNA.

## Introduction

Guanine quadruplexes (G4s) are non-canonical nucleic acid arrangements, whose interest has recently spurred due to their multiple and essential biological functions.^[1]^ G4s are constituted by repeating unit of four guanine bases coupled via Hoogsteen hydrogen bonds, which forms the so-called tetrads. Tetrads are further stabilized by π-stacking, and the high negative electrostatic density is compensated by monovalent cations (usually K^+^ or Na^+^) occupying the central channel.^[2]^ The orientation of the strand connecting the tetrads also gives rise to different arrangements, namely producing parallel, antiparallel, and hybrid topologies.^[3]^ From a biological point of view G4 can be observed both in DNA and RNA sequences (in the following referred to as rG4s), although the latter are much less commonly characterized.^[4]^ This is also probably due to the inherent more pronounced instability of RNA sequences, which complicates their precise structural resolution.^[5]^

G4s were first discovered in the telomeric caps of *Oxytricha nova* and human genomes, with their structures being reported in these contexts, respectively.^[6–10]^ Their capacity to inhibit telomerase^[11]^ contributes to the regulation of cell cycle and the entry in senescence, avoiding the emergence of the immortality phenotype typical of cancer cells. However, G4s have also been evidenced in genes, including oncogene promoters where they participate to their transcriptional regulation.^[10,12]^ While usually the presence of stable and persistent G4s is correlated with gene silencing,^[10]^ in some instances, the presence of G4 structures stimulates transcription.^[13–15]^

Recently, the presence of G4s has also been identified in viral genomes, including both DNA viruses, retroviruses, such as human immunodeficiency virus (HIV),^[16–18]^ and RNA viruses, like SARS-CoV-2,^[5,19–21]^ Zika,^[22]^ and other flaviviruses, making them also ideal targets for novel antiviral therapies.^[23,24]^ Furthermore, the presence of G4s in RNA viral genomes, also proves the importance of these non-canonical arrangement in the framework of RNA regulation.^[25]^ For instance, it has been shown that the translation of VEGF messenger RNA (mRNA) is increased via the interaction of an RNA-binding protein with the rG4, favoring the mRNA targeting to the ribosome.^[26,27]^

Recently, a putative rG4 sequence (PQS) has also been observed in the mRNA transcript of the transmembrane serine protease-2 (TMPRSS2).^[28]^ The localization of this G4 in the 3’ Untranslated region (UTR) of the mRNA could be related to its stabilization and thus to the control of its translation into protein.^[29]^ TMPRSS2 is a membrane-anchored serine protease,^[30]^ mainly expressed by endothelial cells in respiratory and digestive tracts, which is involved in many diverse fundamental biological functions, such as the maintenance of iron homeostasis or the development of idiopathic pulmonary fibrosis.^[31,32]^ Furthermore, TMPRSS2 is also overexpressed in various cancer cells, notably in the case of prostate and breast cancers.^[33]^

For instance, in prostate cancers *TMPRSS2* promoter fuses with the coding sequence of the protooncogene ETS-related gene (ERG) allowing its androgen dependent expression, which promote cancer progression and metastasis development. TMPRSS2 overexpression has also been related to other side effects, and notably to the chronic pain experienced by cancer patients. Because of its involvement in cancer progression, different covalent and non-covalent TMPRSS2 inhibitors have been proposed aimed at poisoning its serine active site.^[34,35]^ The chemical mechanisms underlying the covalent inhibition of TMPRSS2 have also been recently explored using multiscale modelling and free energy methods.^[34]^

More recently, the accessory role of TMPRSS2 in promoting cellular infection by coronaviruses, including SARS-CoV and SARS-CoV-2, has been pointed out.^[28,36–40]^ Indeed, while the recognition mechanism of coronaviruses involves the interaction of the viral Spike protein with the ACE2 cellular receptor,^[41–43]^ TMPRSS2 proteolytic cleavage of SARS-CoV-2 Spike protein favors the fusion of the viral and cellular membrane, hence promoting the infection.^[39,44]^ As a matter of fact, the important role of TMPRSS2 in promoting SARS-CoV-2 infection is also manifested by the fact that this protein has been proposed as a biomarker for COVID-19 possible severe outcomes,^[45]^ while its inhibition has been proposed as an antiviral strategy against SARS-CoV-2.^[28,40]^ The accessory role of TMPRSS2 in promoting viral membrane fusion has also been evidenced for other respiratory viruses, which are also susceptible to generate strong epidemic events, such as influenza A and B viruses.^[46]^ In addition, the important localization of TMPRSS2 in intestinal epithelial cells also suggests its possible involvement in the proteolytic cleavage, favoring the infection of digestive system-targeting viruses.^[47]^

Therefore, stabilizing the mRNA G4 in the 3’ TMPRSS2 UTR represents a suitable original strategy to hamper the (over)expression of the serine protease and thus prevent the spread of respiratory virus infections. Recently, it has been shown that small molecules stabilization of a G4 in the first exon of the TMPRSS2 isoform affects its gene expression and viral replication.^[48]^ In addition, and due to the involvement of TMPRSS2 in cancer progression, the downregulation of its expression by G4 stabilization may also be effective in reducing cancer aggressiveness and metastatic potential.^[48]^ Several molecules have been reported to bind G4s. Common features of promising candidates for interacting with G4 motifs are an extended π-surface area to facilitate stacking atop G-tetrads, a planar geometry, and feature positively charged substituents to enhance interaction with the negatively charged DNA motifs.^[12,49– 55]^ Poor selectivity remains a critical issue with the molecules under investigation. To optimize the molecular design of lead G4 stabilizers we resort to a protocol combining molecular modelling and electronic circular dichroism (ECD) to propose the first predicted structure of the TMPRSS2 rG4.

## Results and Discussion

Contrary to what often reported in the literature, RNA G4s are not necessarily more inclined to form a parallel topology compared to their DNA counterparts. A recent comprehensive analysis of intramolecular G-quadruplex structures revealed that, among the resolved structures to date, there are 33 RNA G4s: fifteen exhibit a parallel topology, nine are antiparallel, five are hybrid, and four are left-handed hybrid.^[56]^ We have recently demonstrated that a RNA guanine-quadruplex structure in the SARS-CoV-2 genome, termed RG-2, can adopt both stable parallel and hybrid G4 conformations, consisting of two rigid tetrads that are nearly ideally π-stacked and connected by short, yet flexible loops.^[5]^ The TMPRSS2 3’ UTR contains a guanine-rich area having the following sequence: 5’-GGGCGGGCGGCCUGCAGGGACAUGGG-3’.

Recently published experimental results have unambiguously shown that this PQS can fold into a G4 arrangement under physiological conditions, adopting a parallel conformation.^[28]^ This conclusion was supported by ECD, FRET melting assays (which we repeated and confirmed, as discussed below), and native PAGE.^[28]^ However, the absence of any experimentally resolved structure of TMPRSS2 G4 has made it necessary to predict its structure at atomic resolution, using a protocol developed by us, and recently employed to predict the structure of two SARS-CoV-2 rG4s.^[5,19]^

This protocol involves, as the first step, the sequence reconstruction by homology modelling.

After the alignment of the TMPRSS2 PQS with different G4-forming sequences, the structure corresponding to the PDB code 2KQH stood out showing the maximum similarity and was, consequently, selected as the initial template. The reconstruction of the sequence first involved the conversion from DNA to RNA and then the addition of the missing nucleotides. Subsequently, two independent molecular dynamics (MD) replicate, each exceeding the µs-time scale, were propagated.

The stability of the arrangement can be also appreciated by the time evolution of the root mean square deviation (RMSD). Indeed, while the tetrad core presents a RMSD of less than 4 Å which remains stable throughout the simulation, the flexible loop shows more pronounced deviations peaking at 8 Å. Unsurprisingly, the total RMSD of the G4 is clearly dominated by the flexible loop as also shown in Figure 1B.

**Figure 1.**
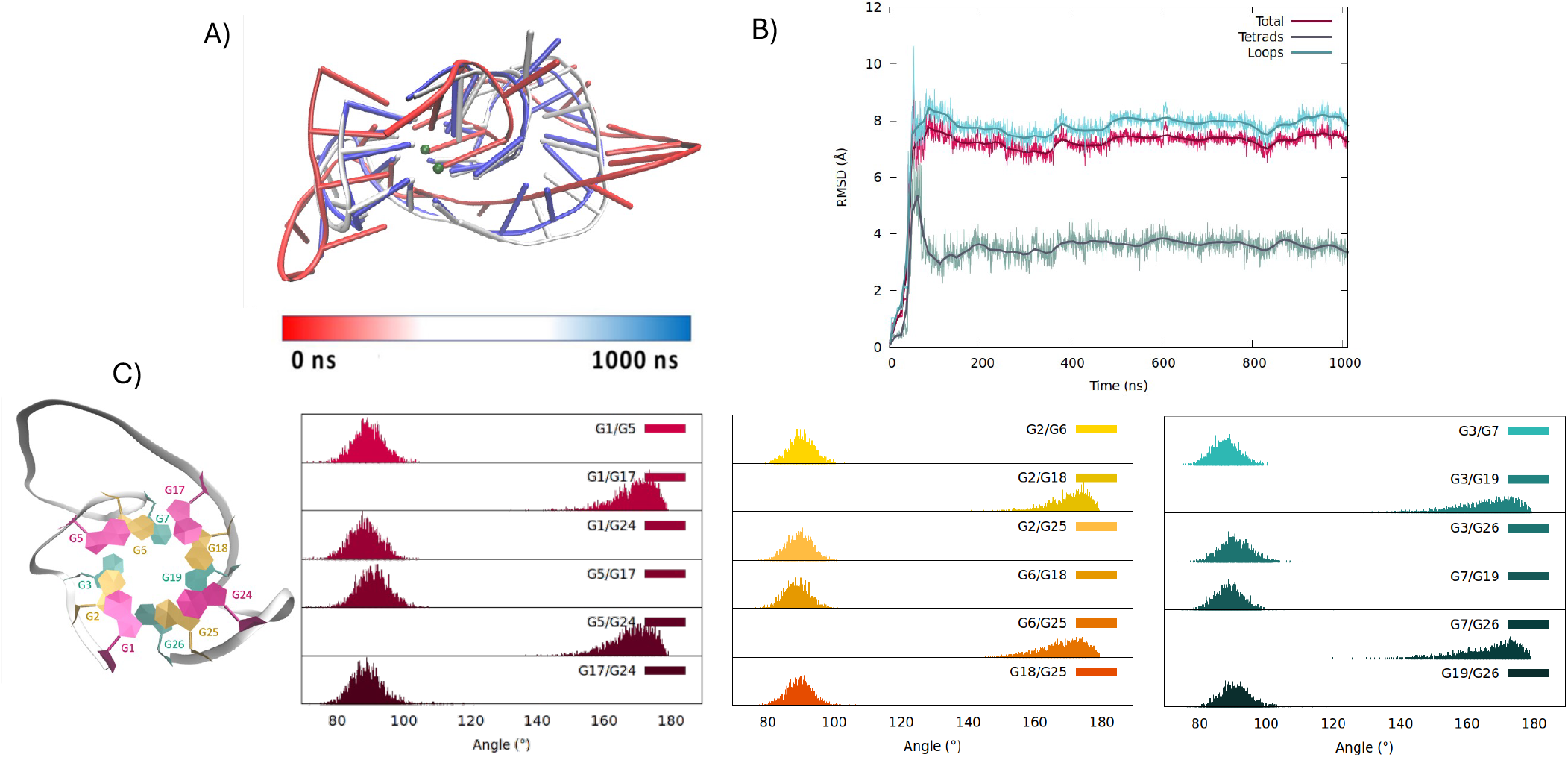
Representative snapshot of the G4 in the TMPRSS mRNA in its parallel arrangement extracted from the MD simulation (A). The corresponding time series of the RMSD for the whole RNA strand, the tetrads and the loops (B). The distribution of the angles between the axis of the guanines composing the first, pink, the second, yellow and the third, forest green, tetrads. The positioning of the different guanine in the RNA structure is also given with the same colour code (C). Analogous results obtained for the second replica are reported in Figure S1.

As shown in Figure 1, the first tetrad is formed by G1, G5, G17, and G24. Conversely, residues G2, G6, G18, and G25 constitute the second tetrad and G3, G7, G19 and G26 originate the last one. From the results presented in Figure 1 we may also see that after a slight structural reorganization observed in the very first steps of the simulation, TMPRSS2 PQS evolves towards a stable parallel G4 arrangement composed of three stacked tetrads and two central K^+^ ions. The tetrads are bridged by three propeller type loops, one of 1 nucleotide (from G3 to G5), a flexible 9 nucleotide long loop (from G7 to G17) and one of 4 nucleotides (G19 to G24). Considering the G4 core, we may observe that the structural parameters extracted from the MD simulations are close to the ideal values both in terms of the twist angle between the tetrads, and the angle between the Hoogsteen-bonded guanines (Figure S3, Table S1-S5).

The global rigidity of the G4 structure is also highlighted by the distance between the centres of mass of each tetrad, which also shows a very limited evolution and oscillation all along the MD simulation (Figure S2).

It is worth noting that TMPRSS2 PQS forms a propeller-type parallel rG4 whit all its guanines adopting an *anti* configuration. Other sequences adopting such a topology tend to form dimeric G4s in which two propeller-type parallel-stranded G4 units are stacked at their 5’ ends.^[57,58]^ However, the analysis of the MD simulation suggests that in our predicted structure, the formation of a dimer would be hampered by the presence of stacked nucleotides on the 5’-3’ tetrads. For instance, in the first replica of the simulation an adenine (A16) belonging to the flexible long loop which stacks on the 5’ tetrad. Similarly, the 3’ prime tetrad has A20 and C8 residues stacking on G19 and G7, respectively. These stacking interactions are stable along all the MD simulation, as demonstrated by time evolution of the distance between the centers of mass of these nucleotides from the guanines belonging to the tetrads (Figure S6). From the graph reported in Figure S6D, nucleotide C8 on top of G7 appears to be less less stable than the other two cases, given the significant difference in distance observed during the simulation.

To support the robustness of our computational protocol and to confirm the G4 arrangement of TMPRSS2, we obtained the simulated ECD spectrum, at the hybrid quantum mechanics/molecular mechanics (QM/MM) level, on top of snapshots issued from the MD simulation and compared it with the experimentally obtained ECD of the same oligomer in K^+^ water solution. The resulting simulated spectrum matches perfectly the shape of the experimental one. Specifically, the presence of a strong positive ellipticity at 260 nm, as well as a negative one at 240 nm align well with previous findings reported and notably supports the parallel conformation of the G4.^[6]^ As it can be appreciated from Figure 2, when considering a global shift of the excitation wavelengths, coherently with our previous work,^[5,19]^ we recover a band shape which is almost totally superposable with the experimental one. Of note, the slight overestimation of the negative peak in the computed spectrum is due to the truncation of the modelled excited states. CD spectra being highly sensitive to even slightly variation in the coupling between the different chromophore, our results clearly support the suitability of the proposed computational-derived structure.

**Figure 2.**
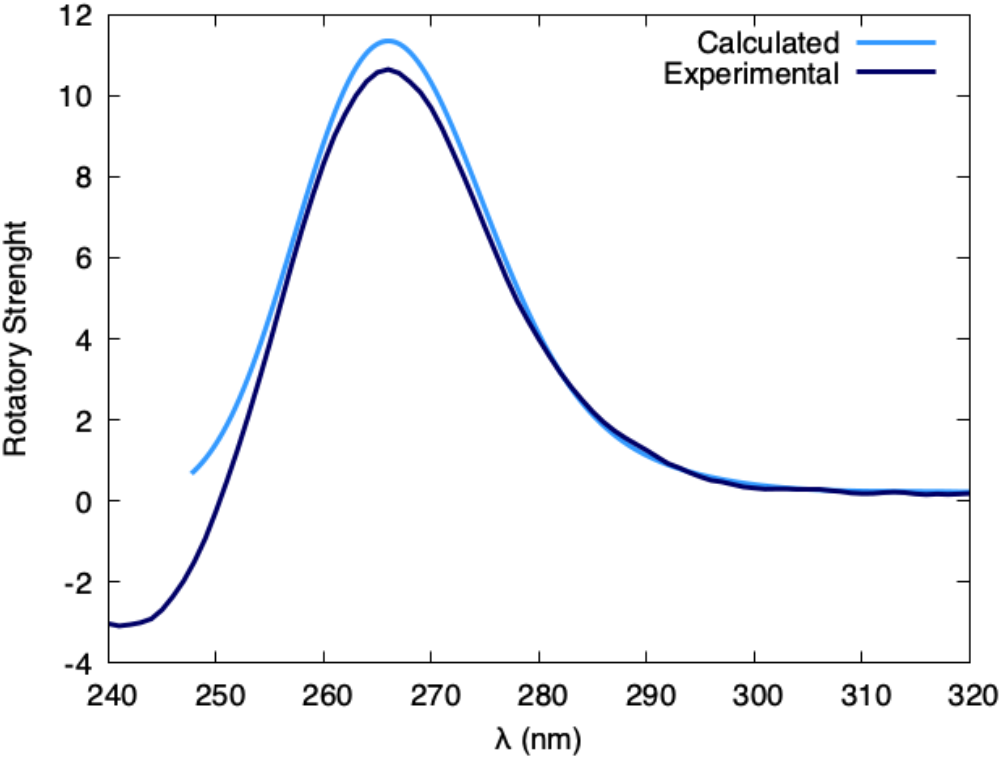
Experimental and modelled ECD spectrum of the TMPRSS2 mRNA G4 in solution. Analogous results obtained for the second replica are reported in the ESI (Figure S7).

Interestingly, the predicted G4 structure of TMPRSS features two long loops, one of which is 9 bases in length. After reviewing the ONQUADRO^[59]^ and G4 database,^[60]^ we did not find any rG4 structure with loops of 9 bases. The traditional theory, although now outdated, suggests that G4 structures typically form with loops of up to 7 nucleotides.^[61]^ it is known that loop length plays a role in G4 stability. Loops of 6 nucleotides or more are generally associated with a destabilizing effect, though this effect is mild. It has also been proposed that no definitive upper limit for loop length can be established for quadruplex formation.^[61]^ We conducted a FRET melting assay to determine the stability of the studied G4. As shown in Figure S4, TMPRSS2 G4 has a Tm of approximately 64 °C, indicating the stability of this structure and its ability to fold under physiological conditions.”

## Conclusions

The biological significance of G4 is nowadays well established, in particular in fundamental nucleic acid sequences related to gene regulation. In this sense, while DNA G4 have been up to date by far the most studied structures, rG4, are nowadays acquiring increased interest. This is also due to the fact that G4 may structure the genome organization of RNA viruses, but also by the fact that they can regulate the translation of mRNA. A PQS sequence has been identified in the mRNA of the TMPRSS2 protein, which can then be tackled to tune its expression. Indeed, controlling the expression level of TMPRSS2 is appealing, since this protease is involved either in cancer progression and in viral infection facilitation. This includes its fundamental role in promoting the entry of SARS-CoV-2 and influenza viruses in the host cell.

Despite the identification of the G4 sequence, its precise structural resolution has remained elusive so far, thus hampering the fine design of potential selective ligands. In this contribution, by leveraging an original protocol developed in our group, and involving the synergic combination of MD simulations and ECD measurements, we have been able to proposed the first structural determination of the G4 in the TMPRSS2 mRNA sequence. We have confirmed that the G4 is forming a rigid core composed of three π-stacked tetrads disposed in a parallel arrangement and flanked by flexible loops. Interestingly, TMPRSS2 is slightly different from other viral RNA viruses, such as RG1 and RG2 in SARS-CoV-2,^[5,19]^ which are composed of only two tetrads. In fact, TMPRSS2 G4 exhibits almost ideal structural parameters and the flexibility is mostly concentrated in the loop regions. Notably the sugar puckering is consistent with the one assumed by RNA sequence, while the dynamics shows the unique population of the 3’-endo conformation. The structural features inferred from the MD simulations are also confirmed by the very nice agreement between the band shape of computed and experimental ECD. Identifying the main structural features of the mRNA G4 will allow us in the future to propose, with the same approach, specific ligands which are suitable to interact with TMPRSS2 G4 and induce its stabilization. This will lead to the development of either original antiviral or oncological drugs, due to the multifaceted role played by TMPRSS2.

## Methodology

### Computational methodology

TMPRSS2 consists of 26 nucleotides that, according to literature, should adopt a parallel G4 structure.^[28]^ To predict at atomic resolution its 3D structure, we used the strategy adopted in recent contributions.^[5,19]^ Specifically, the TMPRSS2 RNA sequence was converted to DNA and the structure of the human telomeric G4 (PDB:1KF1)^[6]^ was used used as a template to force parallel G4 arrangement, enforcing the “Relaxed structure” and “PROSITE” parameters. The best results are reported in Table 1. The best candidate structure to be used as a template for the reconstruction of the TMPRSS2 model was chosen based on a sequence alignment performed using NBCI BLAST.^[62]^ Each of the G4 sequences were aligned with the sequence of TMPRSS2. The sequences of PDB:2KYO^[63]^ and PDB:2KQH,^[64]^ show the best scores and the same nucleotides, have been kept. The choice of the sequence to be used as a template was supported by the evaluation of the structural features using the DSSR G4 database.^[65]^ The first sequence was not selected because, being a dimer and an intermolecular G4, it would not have served as a suitable template. In contrast, the sequence of PDB:2KQH shows high similarity with TMPRSS2. It has three tetrads, parallel arrangement, and it has the best score in terms of similarity.

**Table 1.**
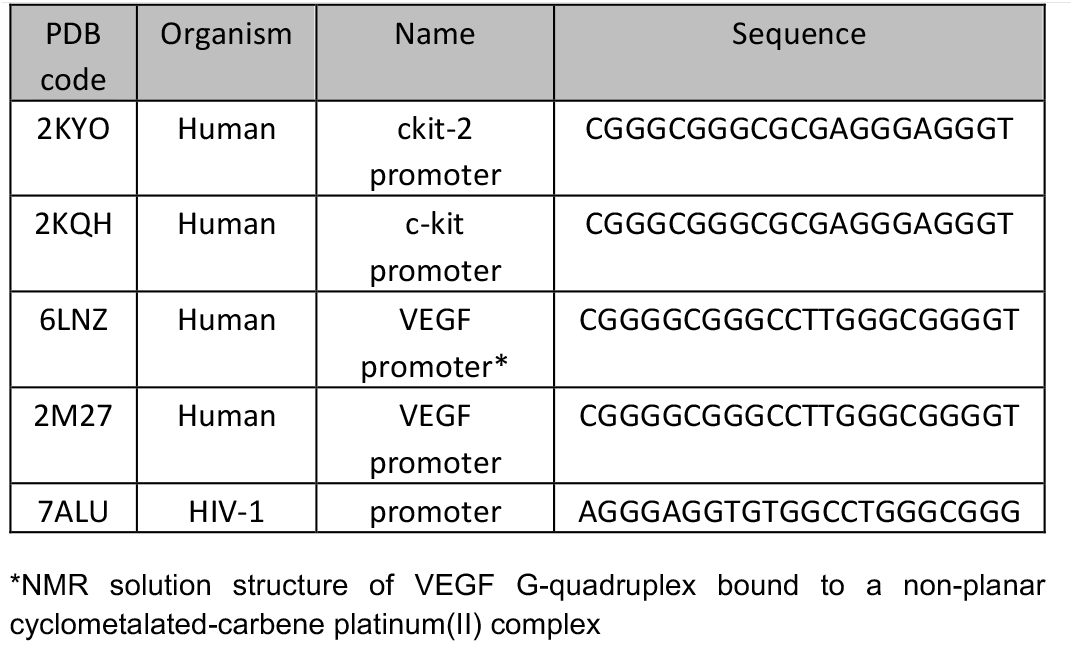
DNA parallel G4 homologous to TMPRSS2 found after searching in the RCSB PDB database.

Construction of the TMPRSS2 rG4 initial models. The parallel TMPRSS model was constructed isolating the equal portion from PDB:2KQH structure and by the reconstructing the missing nucleotides (Figure 3). PyMol (The PyMOL Molecular Graphics System, Version 2.0 Schrödinger, LLC.)^[66]^ was used to convert the DNA structure to RNA and replace all thymine with uracil, using “Mutagenesis Wizard” tool. ChimeraX was used to create the loops and VMD^[67]^ was used to add the central K^+^ cations, necessary for the stability of G4.

**Figure 3.**
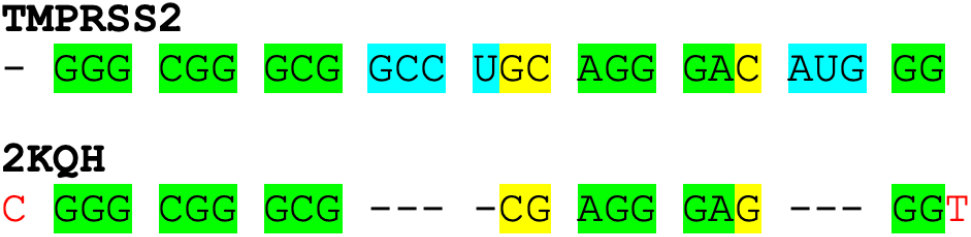
Sequence alignment highlighting the aligned bases (green), the missing nucleotides (blue), the exchanged ones (yellow), the removed bases (red).

### Classical molecular dynamics protocol

The TMPRSS2 initial model is solvated in a TIP3P^[68]^ water box measuring (80 × 80 × 80) Å^3^, with charge neutralization achieved by introducing additional K^+^ potassium ions (7.6*10^−2^ M) via Amber Tleap utilities.^[69]^ TMPRSS2 G4 is modeled using the Amber ff99 force field,^[70]^ augmented with parmbsc0-χOL3 corrections to accurately represent RNA strands.^[71]^ Conformational space exploration is performed through 1 μs molecular dynamics simulations conducted in two replicates using the NAMD code,^[72,73]^ with a time step of 4 fs to integrate Newton’s equations of motion. This choice is enabled by the utilization of the Rattle and Shake algorithm in conjunction with the hydrogen mass repartitioning^[74]^ scheme. Langevin isotherm thermostat^[75]^ and piston^[76]^ are used to maintain an isothermal and isobaric (NPT) ensemble at 300 K and 1 atm pressure. Electrostatic interactions, computed using Particle Mesh Ewald^[77]^ with a cutoff at 9 Å, are incorporated. The system undergoes 1000 steps of minimization, followed by equilibration and thermalization achieved by gradually releasing positional harmonic constraints on heavy atoms over 36 ns. Finally, structural parameters of the nucleic acid G4 are assessed using the script developed by Tsvetkov et al. ^[78]^ et al. and visualized with VMD software. Some structural parameters were obtained using ASC-G4 or WebTetrado webservers.^[79,80]^

### Simulation of the ECD spectra by QM/MM

The protocol used for the simulation to obtain the ECD spectrum is the same to the one used in the previous contributions.^[5,19]^ One hundred snapshots were randomly selected from each MD trajectory, with the eight guanines constituting the two G-quadruplex tetrads and the central K^+^ ion included in the QM partition (Figure S5). Orca software^[81,82]^ was employed to compute the vertical transitions of each snapshot using time-dependent density functional theory (TD-DFT) with the M06-2X functional^[83]^ and the 6-31G basis set. Amber software allowed to perform QM/MM electrostatic embedding,^[84]^ with dangling bonds between MM and QM partitions treated using the link atom approach. The final spectrum was generated by averaging each excitation energy and rotatory strength for all the snapshots. The obtained average vertical transitions have been convoluted with Gaussian functions of Fixed Width at Half Maximum (FWHM) set to 0.4 eV. To match the experimental wavelength a global shift of 37 nm has been consistently applied. Such a shift may help correcting the errors due to the basis set incompleteness, as well as the lack of a polarizable environment. However, we should underline that to support the structural determination we are mostly interested to the recovery of the band shape rather than the absolute excitation energies. As a matter of fact, similar shifts have been applied by us in previous work.^[5,19]^ and are not uncommon in the QM/MM simulations of complex multichromophoric arrangements.

### Experimental ECD

ECD spectra were recorded on a Jasco J-715 spectropolarimeter at 25°C. The parameters were the following: range 400-220 nm, response: 0.5s, accumulation: 4, speed 200 nm/min. For this experiment we used the TMPRSS2 sequence 5’-GGGCGGGCGGCCUGCAGGGACAUGGG-3’ (IDT Technologies). The ODN was resuspended in Tris-KCl buffer (50 mM Tris-HCl, 100 mM KCl, pH 7.4) to yield a 100 μM stock solution expressed in strand units. G-quadruplex (G4) structure was obtainded by heating at 85 °C for 5 minutes, followed by gradual cooling to room temperature.

### FRET Methodology

FRET experiment was performed on a 96-well format Applied BiosystemsTM QuantStudio 6 PCR cycler with a FAM (6-carboxyfluorescein) filter. The sequence of TMPRSS2 (5’-GGGCGGGCGGCCUGCAGGGACAUGGG-3), was modified with FAM and TAMRA (6-carboxy-tetramethylrhodamine) probes at the 5’ and -3’ ends (IDT, Integrated DNA Technologies). Stock solution was prepared solubilizing the lyophilized sequences in RNAase free buffer (Merck). The G4 folding was obtained heating the solutions to 85 °C for 5 min in 60 mM potassium cacodylate buffer (pH 7.4), followed by slowly cooling to room temperature overnight. In the final solution, G4 final concentration was set to 0.2 μM (total volume of 30 μl). Emission data were normalized from 0 to 1.

## Acknowledgements

The authors thank GENCI and Explor computing centers and the Platform P3MB for computational resources. A.M. thanks ANR and CGI for their financial support of this work through Labex SEAM ANR 11 LABX 086, ANR 11 IDEX 05 02. The support of the IdEx “Université Paris 2019” ANR-18-IDEX-0001. A.T. A.S., L.D. and G.B. thank FFR 2021-2022. This work was also financed by European Union – NextGenerationEU – fondi MUR D.M. 737/2021 – project PRJ-0989 (A.T.). A.T. also thanks FFR-D15-162636 project.

## Entry for the Table of Contents

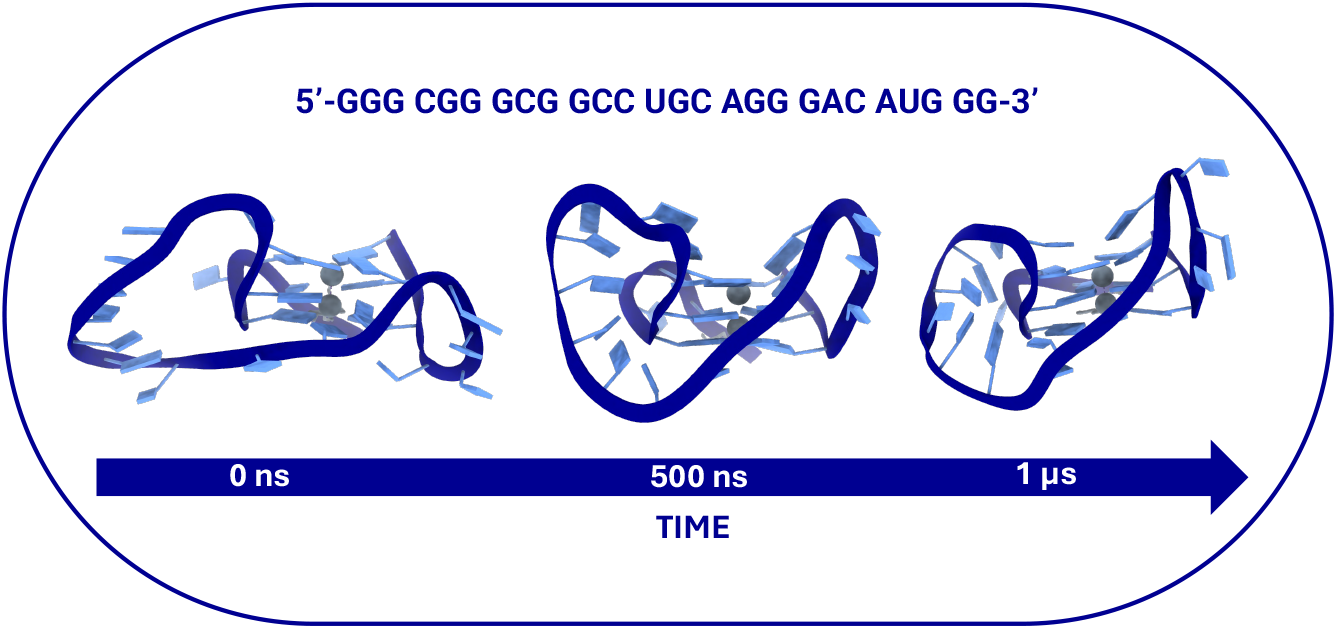

We have determined the structure of a guanine quadruplex motif present in the messenger RNA of the TMPRSS2 protein. By using a combination of molecular dynamics simulations and circular diuchroism spectroscopy we confirm a parallel arrangement involving a rigid tetrad core and flexible loops.

Institute and/or researcher Twitter usernames: @ItodysC, @BIP_BioInorgPa, @LABO_CRAN

## References

[1] R. Hänsel-Hertsch, M. Di Antonio, S. Balasubramanian, Nat Rev Mol Cell Biol 2017, 18, 279–284.

[2] F. Zaccaria, G. Paragi, C. Fonseca Guerra, Physical Chemistry Chemical Physics 2016, 18, 20895–20904.

[3] M. Vorlíčkova, I. Kejnovská, J. Sagi, D. Renčiuk, K. Bednářová, J. Motlová, J. Kypr, Methods 2012, 57, 64–75.

[4] D. Varshney, J. Spiegel, K. Zyner, D. Tannahill, S. Balasubramanian, Nat Rev Mol Cell Biol 2020, 21, 459–474.

[5] L. D’Anna, T. Miclot, E. Bignon, U. Perricone, G. Barone, A. Monari, A. Terenzi, Chem Sci 2023, 14, 11332–11339.

[6] G. Parkinson, M. Lee, S. Neidle, Nature 2002, 417, 876–880.

[7] S. Neidle, FEBS Journal 2010, 277, 1118–1125.

[8] H. Q. Yu, D. Miyoshi, N. Sugimoto, J Am Chem Soc 2006, 128, 15461–15468.

[9] M. P. Horvath, S. C. Schultz, J Mol Biol 2001, 310, 367–377.

[10] S. Balasubramanian, L. H. Hurley, S. Neidle, Nat Rev Drug Discov 2011, 10, 261–275.

[11] A. M. Zahler, J. R. Williamson, T. R. Cech, D. M. Prescott, Nature 1991, 350, 718–720.

[12] A. Froux, L. D’Anna, A. Rainot, C. Neybecker, A. Spinello, R. Bonsignore, R. Rouget, G. Harle, A. Terenzi, A. Monari, S. Grandemange, G. Barone, Inorg Chem Front 2024, DOI 10.1039/D4QI01394H.

[13] S. Lago, M. Nadai, F. M. Cernilogar, M. Kazerani, H. Domíniguez Moreno, G. Schotta, S. N. Richter, Nat Commun 2021, 12, DOI 10.1038/s41467-021-24198-2.

[14] Y. Chen, A. Simeone, L. Melidis, S. M. Cuesta, D. Tannahill, S. Balasubramanian, ACS Chem Biol 2024, 19, 736–742.

[15] C. Ducani, G. Bernardinelli, B. Högberg, B. K. Keppler, A. Terenzi, J Am Chem Soc 2019, 141, 10205–10213.

[16] D. Piekna-Przybylska, M. A. Sullivan, G. Sharma, R. A. Bambara, Biochemistry 2014, 53, 2581–2593.

[17] L. Bonnat, L. Bar, B. Génnaro, H. Bonnet, O. Jarjayes, F. Thomas, J. Dejeu, E. Defrancq, T. Lavergne, Chemistry - A European Journal 2017, 23, 5602–5613.

[18] B. De Nicola, C. J. Lech, B. Heddi, S. Regmi, I. Frasson, R. Perrone, S. N. Richter, A. T. Phan, Nucleic Acids Res 2016, 44, 6442–6451.

[19] T. Miclot, C. Hognon, E. Bignon, A. Terenzi, M. Marazzi, G. Barone, A. Monari, J Phys Chem Lett 2021, 12, 10277–10283.

[20] D. Ji, M. Juhas, C. M. Tsang, C. K. Kwok, Y. Li, Y. Zhang, Brief Bioinform 2021, 22, 1150–1160.

[21] C. Zhao, G. Qin, J. Niu, Z. Wang, C. Wang, J. Ren, X. Qu, Angewandte Chemie - International Edition 2021, 60, 432–438.

[22] A. M. Fleming, Y. Ding, A. Alenko, C. J. Burrows, ACS Infect Dis 2016, 2, 674–681.

[23] S. K. Mukherjee, J. M. Knop, R. Winter, Chemistry - A European Journal 2022, 28, DOI 10.1002/chem.202104182.

[24] G. Qin, C. Zhao, Y. Liu, C. Zhang, G. Yang, J. Yang, Z. Wang, C. Wang, C. Tu, Z. Guo, J. Ren, X. Qu, Cell Discov 2022, 8, DOI 10.1038/s41421-022-00450-x.

[25] E. Ruggiero, S. N. Richter, Bioorg Med Chem Lett 2023, 79, DOI 10.1016/j.bmcl.2022.129085.

[26] K. Niu, X. Zhang, Q. Song, Q. Feng, Int J Mol Sci 2022, 23, 743.

[27] M. J. Morris, Y. Negishi, C. Pazsint, J. D. Schonhoft, S. Basu, J Am Chem Soc 2010, 132, 17831–17839.

[28] G. Liu, W. Du, X. Sang, Q. Tong, Y. Wang, G. Chen, Y. Yuan, L. Jiang, W. Cheng, D. Liu, Y. Tian, X. Fu, Nat Commun 2022, 13, DOI 10.1038/s41467-022-29135-5.

[29] P. Kharel, M. Fay, E. V. Manasova, P. J. Anderson, A. V. Kurkin, J. U. Guo, P. Ivanov, Nat Commun 2023, 14, 205.

[30] L. David K, D. Dongmin, F. Andrea N, H. Markus, S. Brian L, Pain 2015, 156, 923–930.

[31] T. H. Bugge, T. M. Antalis, Q. Wu, Journal of Biological Chemistry 2009, 284, 23177–23181.

[32] H.-H. Li, C.-C. Liu, T.-W. Hsu, J.-H. Lin, J.-W. Hsu, A. F.-Y. Li, Y.-C. Yeh, S.-C. Hung, H.-S. Hsu, Part Fibre Toxicol 2021, 18, 11.

[33] S. Weiss, P. Lamy, M. Rusan, M. Nørgaard, B. P. Ulhøi, M. Knudsen, C. G. Kassentoft, L. Farajzadeh, J. B. Jensen, J. S. Pedersen, M. Borre, K. D. Sørensen, Int J Cancer 2024, 155, 298–313.

[34] A. Spinello, L. D’Anna, E. Bignon, T. Miclot, S. Grandemange, A. Terenzi, G. Barone, F. Barbault, A. Monari, J Phys Chem B 2023, 127, 6287–6295.

[35] C. J. Ko, T. W. Hsu, S. R. Wu, S. W. Lan, T. F. Hsiao, H. Y. Lin, H. H. Lin, H. F. Tu, C. Lee, C. C. Huang, M. J. M. Chen, P. W. Hsiao, H. P. Huang, M. S. Lee, Oncogene 2020, 39, 5950–5963.

[36] L. M. Reinke, M. Spiegel, T. Plegge, A. Hartleib, I. Nehlmeier, S. Gierer, M. Hoffmann, H. Hofmann-Winkler, M. Winkler, S. Pöhlmann, PLoS One 2017, 12, e0179177.

[37] K. Metzdorf, H. Jacobsen, M. C. Greweling-Pils, M. Hoffmann, T. Lüddecke, F. Miller, L. Melcher, A. M. Kempf, I. Nehlmeier, D. Bruder, M. Widera, S. Ciesek, S. Pöhlmann, L. Čičin-Šain, Viruses 2023, 15, 271.

[38] N. Iwata-Yoshikawa, M. Kakizaki, N. Shiwa-Sudo, T. Okura, M. Tahara, S. Fukushi, K. Maeda, M. Kawase, H. Asanuma, Y. Tomita, I. Takayama, S. Matsuyama, K. Shirato, T. Suzuki, N. Nagata, M. Takeda, Nat Commun 2022, 13, 6100.

[39] B. J. Fraser, S. Beldar, A. Seitova, A. Hutchinson, D. Mannar, Y. Li, D. Kwon, R. Tan, R. P. Wilson, K. Leopold, S. Subramaniam, L. Halabelian, C. H. Arrowsmith, F. Bénard, Nat Chem Biol 2022, 18, 963–971.

[40] M. Hoffmann, H. Kleine-Weber, S. Schroeder, N. Krüger, T. Herrler, S. Erichsen, T. S. Schiergens, G. Herrler, N. H. Wu, A. Nitsche, M. A. Müller, C. Drosten, S. Pöhlmann, Cell 2020, 181, 271–280.

[41] Q. Wang, Y. Zhang, L. Wu, S. Niu, C. Song, Z. Zhang, G. Lu, C. Qiao, Y. Hu, K. Y. Yuen, Q. Wang, H. Zhou, J. Yan, J. Qi, Cell 2020, 181, 894-904.e9.

[42] E. S. Brielle, D. Schneidman-Duhovny, M. Linial, Viruses 2020, 12, 497.

[43] R. Yan, Y. Zhang, Y. Li, L. Xia, Y. Guo, Q. Zhou, Science (1979) 2020, 367, 1444–1448.

[44] D. Bestle, M. R. Heindl, H. Limburg, T. van Lam van, O. Pilgram, H. Moulton, D. A. Stein, K. Hardes, M. Eickmann, O. Dolnik, C. Rohde, H. D. Klenk, W. Garten, T. Steinmetzer, E. Böttcher-Friebertshäuser, Life Sci Alliance 2020, 3, e202000786.

[45] J. D. Strope, C. H. C. PharmD, W. D. Figg, The Journal of Clinical Pharmacology 2020, 60, 801–807.

[46] H. Limburg, A. Harbig, D. Bestle, D. A. Stein, H. M. Moulton, J. Jaeger, H. Janga, K. Hardes, J. Koepke, L. Schulte, A. R. Koczulla, B. Schmeck, H.-D. Klenk, E. Böttcher-Friebertshäuser, J Virol 2019, 93, e00649.

[47] M. Esumi, M. Ishibashi, H. Yamaguchi, S. Nakajima, Y. Tai, S. Kikuta, M. Sugitani, T. Takayama, M. Tahara, M. Takeda, T. Wakita, Hepatology 2015, 61, 437–446.

[48] A. De Magis, P. Schult, A. Schönleber, R. Linke, K. U. Ludwig, B. M. Kümmerer, K. Paeschke, BMC Biol 2024, 22, 5.

[49] E. Ruggiero, S. N. Richter, Nucleic Acids Res 2018, 46, 3270–3283.

[50] A. Terenzi, H. Gattuso, A. Spinello, B. K. Keppler, C. Chipot, F. Dehez, G. Barone, A. Monari, Antioxidants 2019, 8, DOI 10.3390/antiox8100472.

[51] R. Bonsignore, F. Russo, A. Terenzi, A. Spinello, A. Lauria, G. Gennaro, A. M. Almerico, B. K. Keppler, G. Barone, J Inorg Biochem 2018, 178, 106–114.

[52] A. Terenzi, R. Bonsignore, A. Spinello, C. Gentile, A. Martorana, C. Ducani, B. Högberg, A. M. Almerico, A. Lauria, G. Barone, RSC Adv. 2014, 4, 33245–33256.

[53] L. D’Anna, S. Rubino, C. Pipitone, G. Serio, C. Gentile, A. Palumbo Piccionello, F. Giannici, G. Barone, A. Terenzi, Dalton Transactions 2023, 52, 2966–2975.

[54] W. Streciwilk, A. Terenzi, R. Misgeld, C. Frias, P. G. Jones, A. Prokop, B. K. Keppler, I. Ott, ChemMedChem 2017, 12, 214–225.

[55] J. Figueiredo, J.-L. Mergny, C. Cruz, Life Sci 2024, 340, 122481.

[56] M. Farag, L. Mouawad, Nucleic Acids Res 2024, 52, 3522–3546.

[57] N. Q. Do, K. W. Lim, M. H. Teo, B. Heddi, A. T. Phan, Nucleic Acids Res 2011, 39, 9448–9457.

[58] N. Q. Do, A. T. Phan, Chemistry – A European Journal 2012, 18, 14752–14759.

[59] T. Zok, N. Kraszewska, J. Miskiewicz, P. Pielacinska, M. Zurkowski, M. Szachniuk, Nucleic Acids Res 2022, 50, D253–D258.

[60] X.-J. Lu, Nucleic Acids Res 2020, DOI 10.1093/nar/gkaa426.

[61] A. Guédin, J. Gros, P. Alberti, J.-L. Mergny, Nucleic Acids Res 2010, 38, 7858–7868.

[62] J. Mark, Z. Irena, R. Yan, M. Yuri, M. Scott, T. L. Madden, Nucleic Acids Res 2008, 36, W5–W9.

[63] V. Kuryavyi, A. T. Phan, D. J. Patel, Nucleic Acids Res 2010, 38, 6757–6773.

[64] S.-T. D. Hsu, P. Varnai, A. Bugaut, A. P. Reszka, S. Neidle, S. Balasubramanian, J Am Chem Soc 2009, 131, 13399–13409.

[65] X. J. Lu, H. J. Bussemaker, W. K. Olson, Nucleic Acids Res 2015, 43, e142.

[66] Schrödinger LLC, The PyMOL Molecular Graphics System, Version∼1.8, 2015.

[67] W. Humphrey, A. Dalke, K. Schulten, J Mol Graph 1996, 14, 33–38.

[68] P. Mark, L. Nilsson, Journal of Physical Chemistry A 2001, 105, 9954–9960.

[69] D. A. Case, J. T. Berryman, R. M. Betz, D. S. Cerutti, I. I. I. T. E. Cheatham, T. A. Darden, R. E. Duke, T. J. Giese, H. Gohlke, A. W. Goetz, N. Homeyer, S. Izadi, P. Janowski, J. Kaus, A. Kovalenko, T. S. Lee, S. Le Grand, P. L. T. Luchko, R. Luo, d K. M. M. B. Madej a, G. Monard, P. Needham, H. Nguyen, H. T. Nguyen, I. Omelyan, A. Onufriev, D. R. Roe, A. Roitberg, R. Salomon-Ferrer, C. L. Simmerling, W. Smith, J. Swails, R. C. Walker, J. Wang, R. M. Wolf, X. Wu, D. M. York, P. A. Kollman, 2022, Amber 2022, University of California, San Francisc.

[70] K. Lindorff-Larsen, S. Piana, K. Palmo, P. Maragakis, J. L. Klepeis, R. O. Dror, D. E. Shaw, Proteins: Structure, Function, and Bioinformatics 2010, 78, NA-NA.

[71] M. Zgarbová, M. Otyepka, J. Šponer, A. Mládek, P. Banáš, T. E. Cheatham, P. Jurečka, J Chem Theory Comput 2011, 7, 2886–2902.

[72] J. C. Phillips, R. Braun, W. Wang, J. Gumbart, E. Tajkhorshid, E. Villa, C. Chipot, R. D. Skeel, L. Kalé, K. Schulten, J Comput Chem 2005, 26, 1781–1802.

[73] J. C. Phillips, D. J. Hardy, J. D. C. Maia, J. E. Stone, J. V. Ribeiro, R. C. Bernardi, R. Buch, G. Fiorin, J. Hénin, W. Jiang, R. McGreevy, M. C. R. Melo, B. K. Radak, R. D. Skeel, A. Singharoy, Y. Wang, B. Roux, A. Aksimentiev, Z. Luthey-Schulten, L. V. Kalé, K. Schulten, C. Chipot, E. Tajkhorshid, Journal of Chemical Physics 2020, 153, 044130.

[74] C. W. Hopkins, S. Le Grand, R. C. Walker, A. E. Roitberg, J Chem Theory Comput 2015, 11, 1864–1874.

[75] R. L. Davidchack, R. Handel, M. V. Tretyakov, Journal of Chemical Physics 2009, 130, 234101.

[76] S. E. Feller, Y. Zhang, R. W. Pastor, B. R. Brooks, J Chem Phys 1995, 103, 4613–4621.

[77] T. Darden, D. York, L. Pedersen, J Chem Phys 1993, 98, 10089–10092.

[78] V. Tsvetkov, G. Pozmogova, A. Varizhuk, J Biomol Struct Dyn 2016, 34, 705–715.

[79] M. Farag, C. Messaoudi, L. Mouawad, Nucleic Acids Res 2023, 51, 2087–2107.

[80] B. Adamczyk, M. Zurkowski, M. Szachniuk, T. Zok, Nucleic Acids Res 2023, 51, W607–W612.

[81] F. Neese, Wiley Interdiscip Rev Comput Mol Sci 2012, 2, 73–78.

[82] F. Neese, F. Wennmohs, U. Becker, C. Riplinger, Journal of Chemical Physics 2020, 152, 224108.

[83] Y. Zhao, D. G. Truhlar, Theor Chem Acc 2008, 120, 215–241.

[84] A. W. Götz, M. A. Clark, R. C. Walker, J Comput Chem 2014, 35, 95–108.

