## Supplementary Information for "Resolving the Structure of a Guanine Quadruplex in TMPRSS2 Messenger RNA by Circular Dichroism and Molecular Modeling"

a) Department of Biological, Chemical and Pharmaceutical Sciences, University of Palermo, 90126 Palermo, Italy; b) Université de Lorraine and CNRS, UMR 7019 LPCT, F-54000 Nancy, France;c) Université Paris Cité and CNRS, ITODYS F-75006 Paris, France; d) Fondazione Ri.MED, Via Filippo Marini 14, 90128, Palermo, Italy

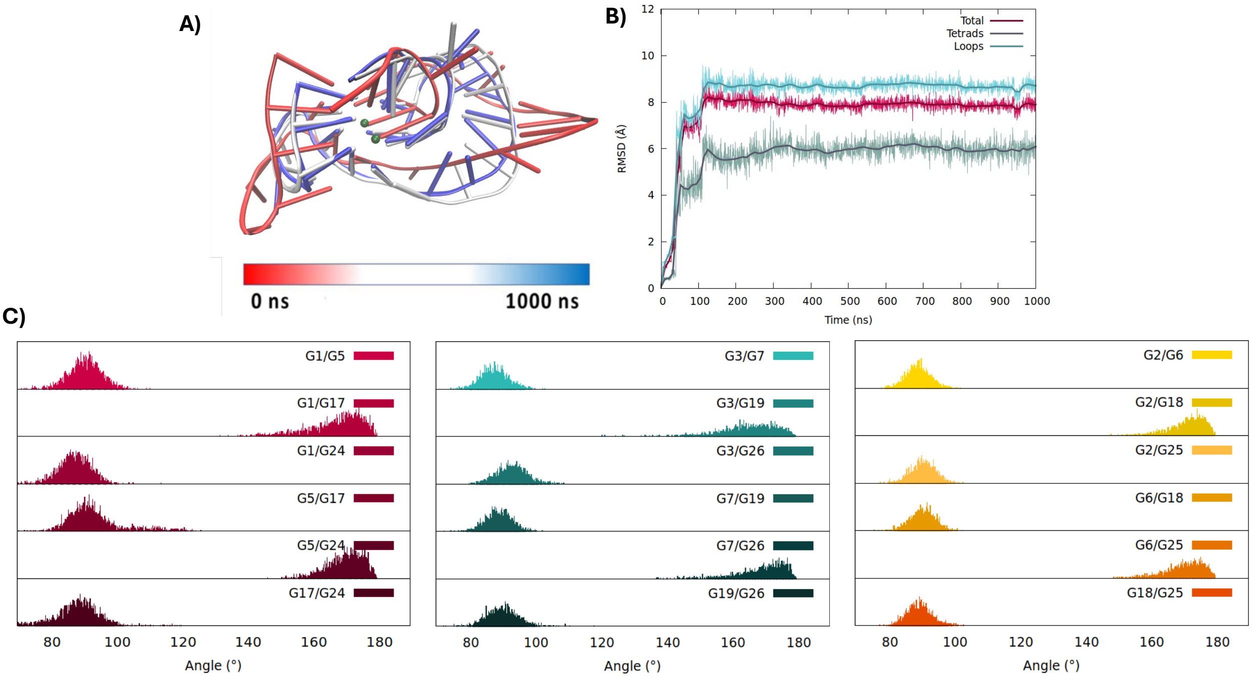


**Figure S1**. Representative snapshot of the G4 in the TMPRSS mRNA in its parallel arrangement extracted from the second replica of the MD simulation (**A**). The corresponding time series of the RMSD for the whole RNA strand, the tetrads and the loops (**B**). The distribution of the angles between the axis of the guanines composing the first, pink, the second, yellow and the third, forest green, tetrads. The positioning of the different guanine in the RNA structure is also given with the same colour code. (**C**)

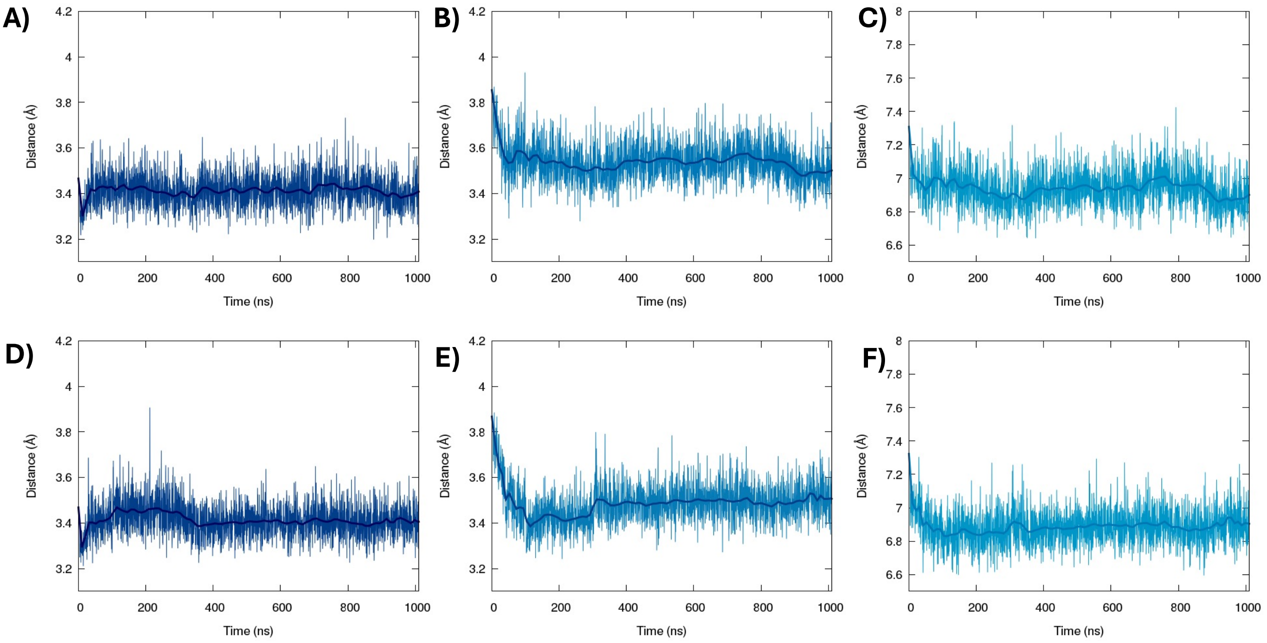


**Figure S2**. Time series of the distance between the first and second tetrad (**A**,**D**), second and third tetrad (**B**,**E**), first and third one (**C**,**F**). The graphs refer to the first (**A**-**C**) and the second replica (**D**-**F**).

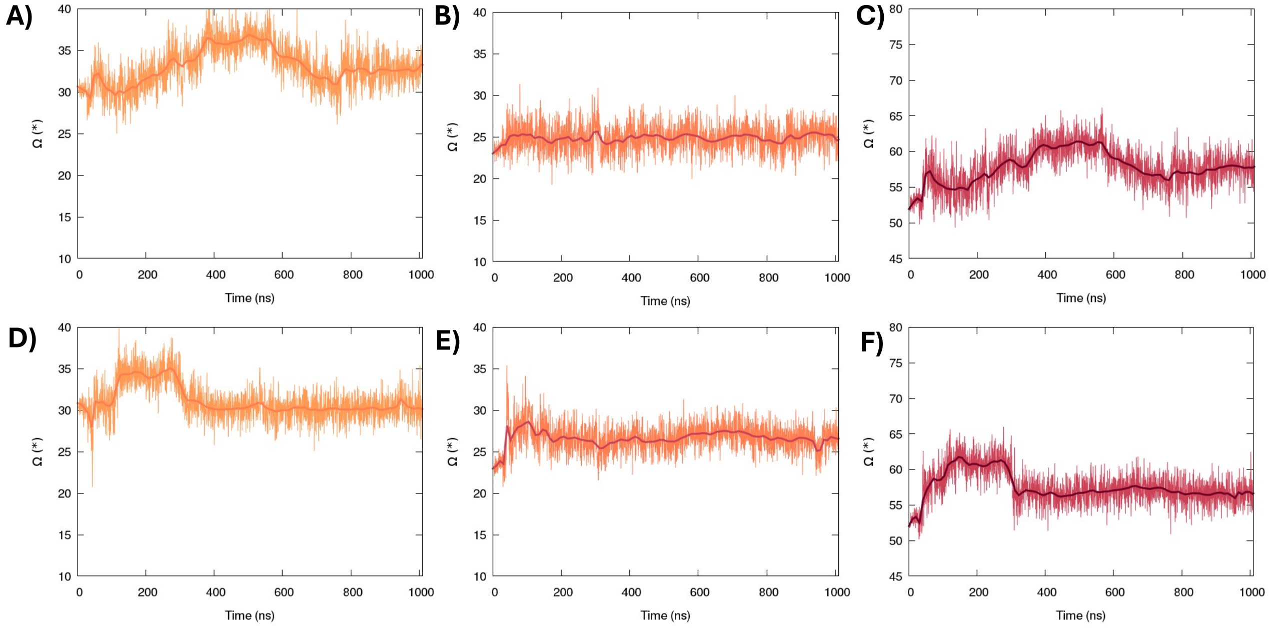


**Figure S3**. Time series of the twist angle (Ω) between the first and the second tetrad (**A**,**D**), for the second and the third tetrad (**B**,**E**), for the first and the third one (**C**,**F**). The graphs refer to the first (**A**-**C**) and the second replica (**D**-**F**).

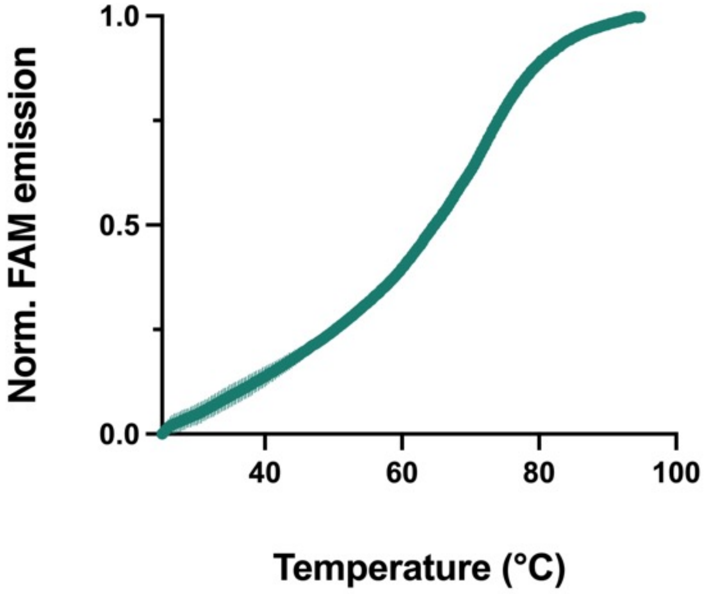


**Figure S4.** FRET melting curve of TMPRSS2 G4 (60 mM potassium cacodylate buffer, pH = 7.4)

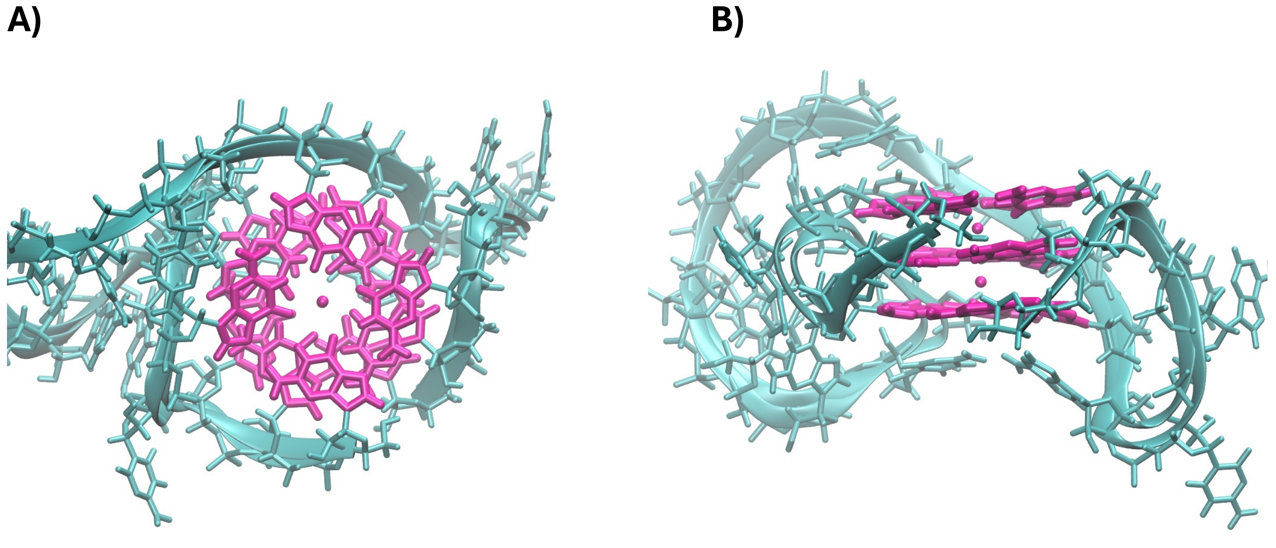


**Figure S5**. Top (A) and side view (B) of the MM (cyan) and QM (magenta) partitions used for the simulation of the ECD spectra.

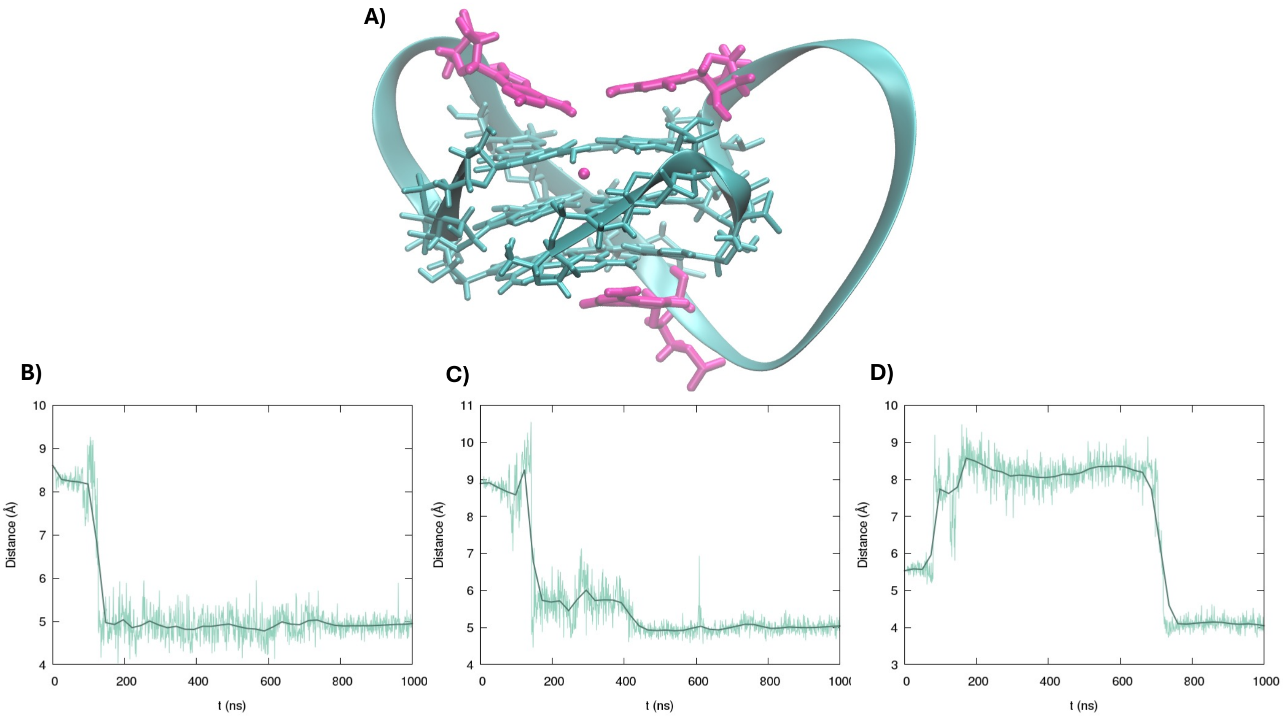


**Figure S6.** Cartoon showing the presence of stacking nucleotides (magenta) on 5'-3' tetrads (A). Time evolution of the distance between the centres of mass of nucleotides A20-G19 (B), A16-G17 (C), C8-G7 (D).

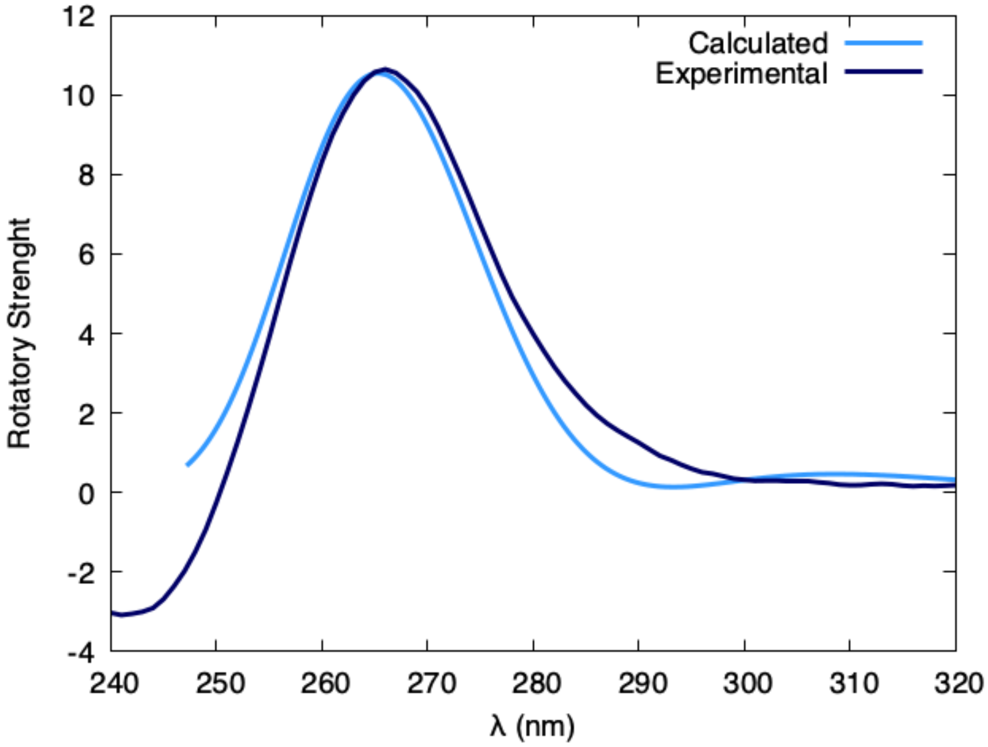


**Figure S7**. Simulated ECD spectra obtained from the second replica compared with the experimental results.

**Table S1.** Twist angle between the tetrad for the first and second replica.

| Twist Angles | 1°-2° tetrad | 2°-3° tetrad | 1°-3° tetrad |
|---|---|---|---|
| First replica | 33.0 ± 2° | 25.1 ± 2° | 57.9 ± 3° |
| Second replica | 31.1 ± 2° | 26.6 ± 2° | 57.5 ± 2° |

**Table S2.** Distance between the tetrads for the first and second replica.

| Distance between quartet | 1°-2° tetrad | 2°-3° tetrad | 1°-3° tetrad |
|---|---|---|---|
| First replica | 3.40 ± 0.07 | 3.53 ± 0.08 | 6.93 ± 0.11 |
| Second replica | 3.41 ± 0.07 | 3.48 ± 0.09 | 6.88 ± 0.10 |

**Table S3.** Distribution of the angles between the axis of the guanines composing the first tetrad for the first and second replica.

| | G1-G5 | G1-G17 | G1-G24 | G5-G17 | G5-G24 | G17-G24 |
|---|---|---|---|---|---|---|
| **First replica** | 90.07 ± 4.34 | 170.02 ± 6.01 | 88.84 ± 4.47 | 91.37 ± 4.76 | 168.92 ± 6.33 | 89.66 ± 5.01 |
| **Second replica** | 90.74 ± 4.80 | 167.78 ± 8.00 | 87.61 ± 4.83 | 93.29 ± 8.13 | 170.29 ± 5.32 | 88.20 ± 6.97 |

**Table S4.** Distribution of the angles between the axis of the guanines composing the second tetrad for the first and second replica.

| | G2-G6 | G2-G18 | G2-G25 | G6-G18 | G6-G25 | G18-G25 |
|---|---|---|---|---|---|---|
| **First replica** | 90.07 ± 4.34 | 170.02 ± 6.01 | 88.84 ± 4.47 | 91.37 ± 4.76 | 168.92 ± 6.33 | 89.66 ± 5.01 |
| **Second replica** | 90.74 ± 4.80 | 167.78 ± 8.00 | 87.61 ± 4.83 | 93.29 ± 8.13 | 170.29 ± 5.32 | 88.20 ± 6.97 |

**Table S5.** Distribution of the angles between the axis of the guanines composing the third tetrad for the first and second replica.

| | G3-G7 | G3-G19 | G3-G26 | G7-G19 | G7-G26 | G19-G26 |
|---|---|---|---|---|---|---|
| **First replica** | 88.45 ± 3.78 | 166.15 ± 8.90 | 91.35 ± 4.32 | 89.39 ± 4.02 | 167.72 ± 8.49 | 91.31 ± 4.27 |
| **Second replica** | 87.96 ± 3.84 | 165.89 ± 7.95 | 93.05 ± 4.34 | 89.04 ± 8.76 | 170.21 ± 6.72 | 90.24 ± 4.43 |
